# Extended “Constant Field” equations: an electrodiffusion model of NMDA receptors

**DOI:** 10.1101/221408

**Authors:** Keivan Moradi, Kamran Moradi, Fatemeh Bakouie, Shahriar Gharibzadeh

## Abstract

Ion channels are experimentally divided into Ohmic and non-Ohmic channels. Similarly, a theoretical dichotomy exists in the flux equations of these channels. The “constant field” or “Goldman-Hodgkin and Katz” current equation is a commonly accepted analytic model of electrodiffusion in channels. This equation, however, seems to have some shortcomings, such as the inability to explain the nonlinear relation between the single channel current and the membrane potential in non-Ohmic channels. In this study, we introduce the “general” and “extended” versions of this equation that are applicable to both Ohmic and non-Ohmic channels. Our results showed the ability of these equations to simulate the NMDA receptor current, as a non-Ohmic channel, not only in the symmetric “uni-ionic” solutions but also in the asymmetric “bi-ionic” solutions of sodium and calcium ions. Our equations reveal for the first time that sodium ions like calcium ions are able to block the NMDA receptors current. It seems that the previously identified cation binding-sites have a prominent role in changing the diffusion coefficient inside the channel. We also propose the concentration-dependent change in the asymmetry of diffusion coefficient as a remarkably effective mechanism for shaping the channel’s *I-V* curve.

## 1. Introduction

The permeation models of ion channels are of particular importance in some computational studies, as in the modeling of synaptic plasticity, e.g. see(Holmes and Levy 1990; Badoual et al. 2006). The most commonly used analytic electrodiffusion equation with a special significance in the calculation of the ionic flux of channels is the “constant field” or the “Goldman-Hodgkin and Katz” current equation(Goldman 1943; Hodgkin et al. 1949). In this equation, the calculation of ionic flux is based on the channel’s permeability, the concentration difference (diffusion potential) and the electric potential. A set of equations have been derived from the constant field current equation in order to calculate the ionic permeability of channels to different ions(Lewis 1979). This equation also has a particular theoretical significance for its ability to predict the Nernst equation in electrical potentials equal to zero, and the Ohm’s law in diffusion potentials equal to zero(Hille 2001). Nevertheless, according to the experimental findings, some channels do not show an Ohmic response in symmetric uni-ionic solutions, where a same concentration of only one ionic species exists in both sides of the channel and as a result, the diffusion potential is equal to zero(Iino et al. 1997; Kandel 2012). At present, the ionic flux of these so-called non-Ohmic channels is calculated by complex thermodynamic models(Iino et al. 1997; Hille 2001) or non-constant field numerical electrodiffusion equations(D. P. Chen 2005; Feng 2003). In other words, currently, no unified analytic electrodiffusion equation of Ohmic and non-Ohmic ion channels exists and this has resulted in a dichotomy between the flux equations for channels.

Moreover, electrodiffusion theory should keep up with the recent advances in experimental techniques that have made it possible to record the electrical activities of the “dendritic spines”(Takahashi et al. 2012). The reason is, having a small surface to volume ratio, the dendritic spines are faced with vast fluctuations in diffusion potential that are comparable in the amounts to the fluctuations of the electric potentials. However, being one of the essential parts of the dendritic spines, NMDA receptor (NMDAR) as an example, is currently modeled with Ohm’s law in the majority of computational studies, e.g. see(Harnett et al. 2012; Grunditz et al. 2008). Yet, there should be three conditions in place for any channel, like NMDARs, to be modeled by Ohm’s law and NMDARs are not subject to these conditions based on the following reasons:

First, the electric potential of the membrane being studied (*V*) should change in a range far beyond the reversal potentials (*E*_rev_) of its channels (e.g. [*V-E_rev_*] >> *RT/F* ≈ 25 mV) so that the diffusional forces in the membrane become negligible compared to the electrical ones. As a result, it would be possible to consider that the current passing through the channel is approximately a linear function of the membrane voltage. In NMADARs, however, the magnesium ions block the channel in the hyperpolarized membrane potentials. Therefore, the NMDARs’ current reaches its peak at about -20 mV, which is close to its *E*_rev_ (≈ 0 mV). In other words, NMDAR current reaches its peak when ([*V*-*E*_rev_]<25 mV).

Second, in the Ohmic model, when the calculation of the ionic fluxes near the *E*_rev_ is required, it is necessary to assume that the channel is approximately permeable to just one ionic species. The reason is that the Ohmic model calculates a zero flux at *E*_rev_, regardless of the concentration and the number of ionic species permeating through the channel. This assumption is valid in “uni-ionic” channels where only one ionic species is permeating the channel. In a “multi-ionic” channel, however, although the net current near the *E*_rev_ is equal to zero, the net flux of each ionic species may or may not be. The NMDARs, also, have a multi-ionic channel that is permeable to a mixture of ions, consisting of Na^+^, K^+^ and Ca^2+^ ions, in physiological solutions. Therefore in these channels, the calculation of the ionic flux of each of these ions is difficult near the *E*_rev_.

Third, by changing the solution concentration, the ionic flux is able to modify both the *E*_rev_ and the conductance of an Ohmic channel. Thus, in order to calculate the ionic flux accurately, these parameters should be updated constantly whenever the calculation of the ionic flux is of particular importance. In a uni-ionic channel, the estimation of the *E*_rev_ is an “equilibrium problem” that is solved by one of the derivatives of the Boltzmann equation, the Nernst equation. In multi-ionic channels like NMDARs, however, this estimation is a “non-equilibrium electrodiffusion problem” that is solved by the “constant field” *voltage* equation, which is derived from the original constant field *current* (oCFc) equation(Hille 2001; De Schutter 2010). Thus, it is more reasonable to use the oCFc equation directly to calculate the current.

Unfortunately, all aspects of ion permeation in NMDARs, could not be explained by the recommended oCFc equation (De Schutter 2010) that is used for the simulation of these receptors (Badoual et al. 2006; Holmes and Levy 1990). According to the following statements, some conditions in NMDARs could not be explained by oCFc equation:

1. Experimental findings have shown that NMDARs are non-Ohmic channels as they have non-linear *I-V* curves in symmetric uni-ionic solutions of Na^+^ ions(Iino et al. 1997).
2. It has been shown in a recent study that the relative permeability value of Ca^2+^ to Na^+^ *(P_ca_/P_Na_)* that is calculated by the oCFc equations is incompatible with the calculated values that are based on the fractional calcium current *(P_f_)* measurements, which is a theory independent method (Jatzke et al. 2002).
3. The external calcium blocks the inward but not the outward permeation of ions in a concentration-dependent manner (Jahr and Stevens 1987, 1993). The oCFc equation, however, is unable to model this type of block. The reason is that although the oCFc equation can be rewritten so that it can calculate the inward and the outward currents separately (see Eq. 17), it is not possible to apply a calcium block coefficient to just the inward limb of this equation. The reason is, by doing so, some of the underlying assumptions of the oCFc equation that the ions have an equal chance to enter the channel’s mouth on each side, and that the diffusion coefficient of ions have a constant value throughout the channel, are violated.

### Previous models of the NMDAR current

We are aware of only two studies that have tried to formulate the experimentally recorded NMDARs current in a single channel mode. One of them, which is based on Ohm’s law, considers different conductances for the inward and the outward currents(Jahr and Stevens 1993). In this model, the channel conductance results from the weighted sum of the inward and outward conductances. The *E*_rev_ calculation is based on a special form of the Nernst equation. Apart from the discussed problems of Ohmic models, one problem with this model, to our knowledge, is that its *E*_rev_ equation does not have an analytical solution. Moreover, the authors have approached the problem from a misleading theoretical standpoint since the calculation of *E*_rev_ in multi-ion channels in mixed ionic solutions is a non-equilibrium problem that is incompatible with Nernst’s equilibrium equation.

In the second study, Iino et al. have used a thermodynamic model that is derived from the biophysically criticized Eyring-Läuger theory of the ion permeation in channels (Cooper et al. 1988), to simulate the current permeation of NMDARs in a single channel mode(Iino et al. 1997). They consider a “one-ion, three-barrier” model for the NMDA channels. Their experimental data seem to be more precise than those of Jahr and Stevens are, because their experimental data is limited to the main conductance states of the NMDAR current. In some parts of Jahr and Stevens study, however, the main and sub-conductance states are mixed together; while, in another study, it has been shown that the sub-conductance states have a different *I-V* curve than those of the main states(Schneggenburger and Ascher 1997). Consequently, the sub-conductance states may have a different permeability to permeant ions. Despite being insightful, we believe that the computational parts of Iino et al. study have some inaccuracies. In the Fig. 2 of that study, for example, they have reported a parameter named 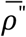 that is not used in the calculations of the sub-functions of their current equation (i.e. *R_Na_*(*ϕ*) or *S_Na_*(*ϕ*)). Their equations also have a significant difference with what was originally introduced by Chang et al. (Chang et al. 1994). Without further optimizations also, we were not able to simulate their experimental results with the estimated parameters and the reported equations of their study (and even with that of Chang et al. study).

**Fig. 1.**
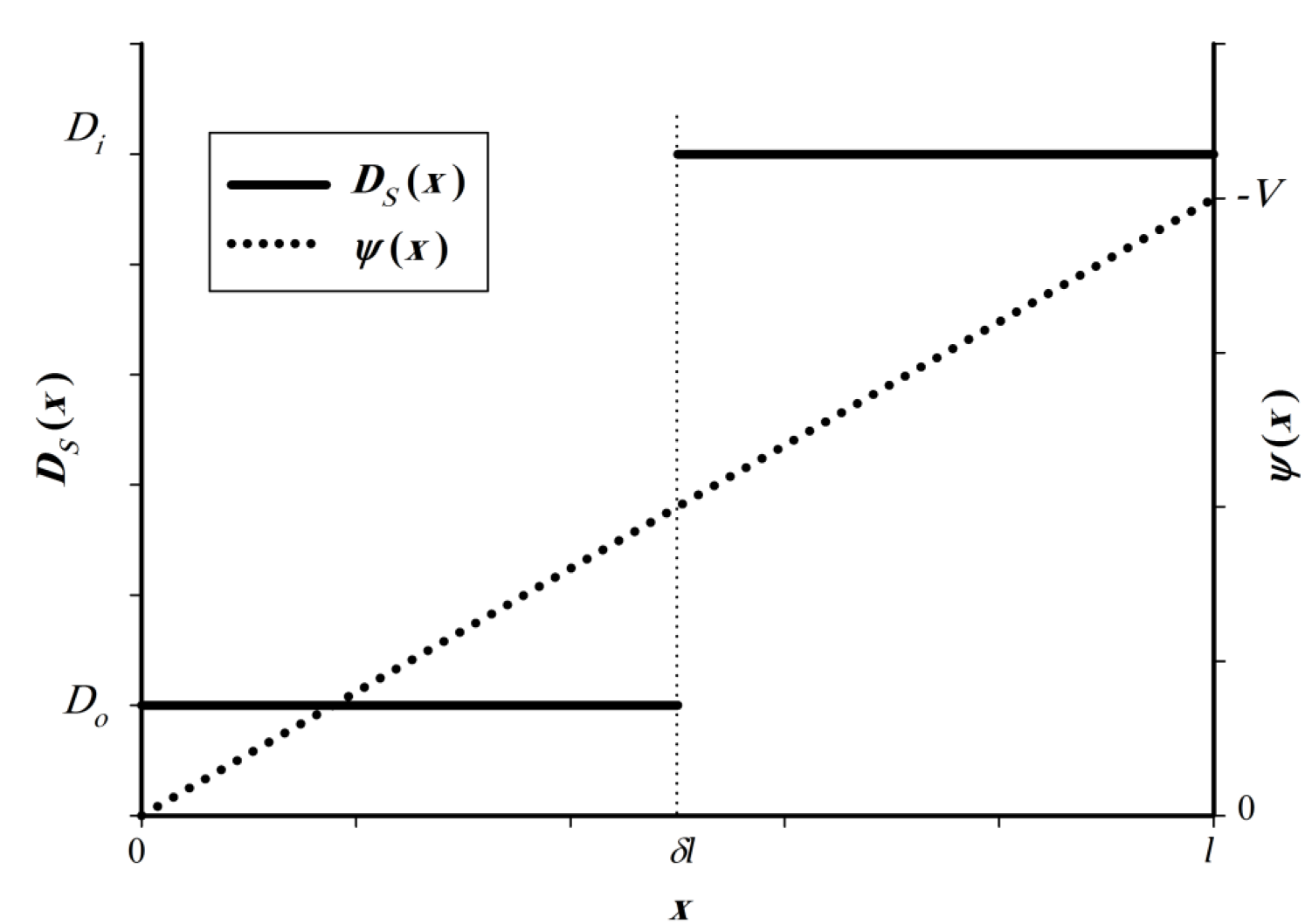
The model’s parameters within the channel.

**Fig. 2.**
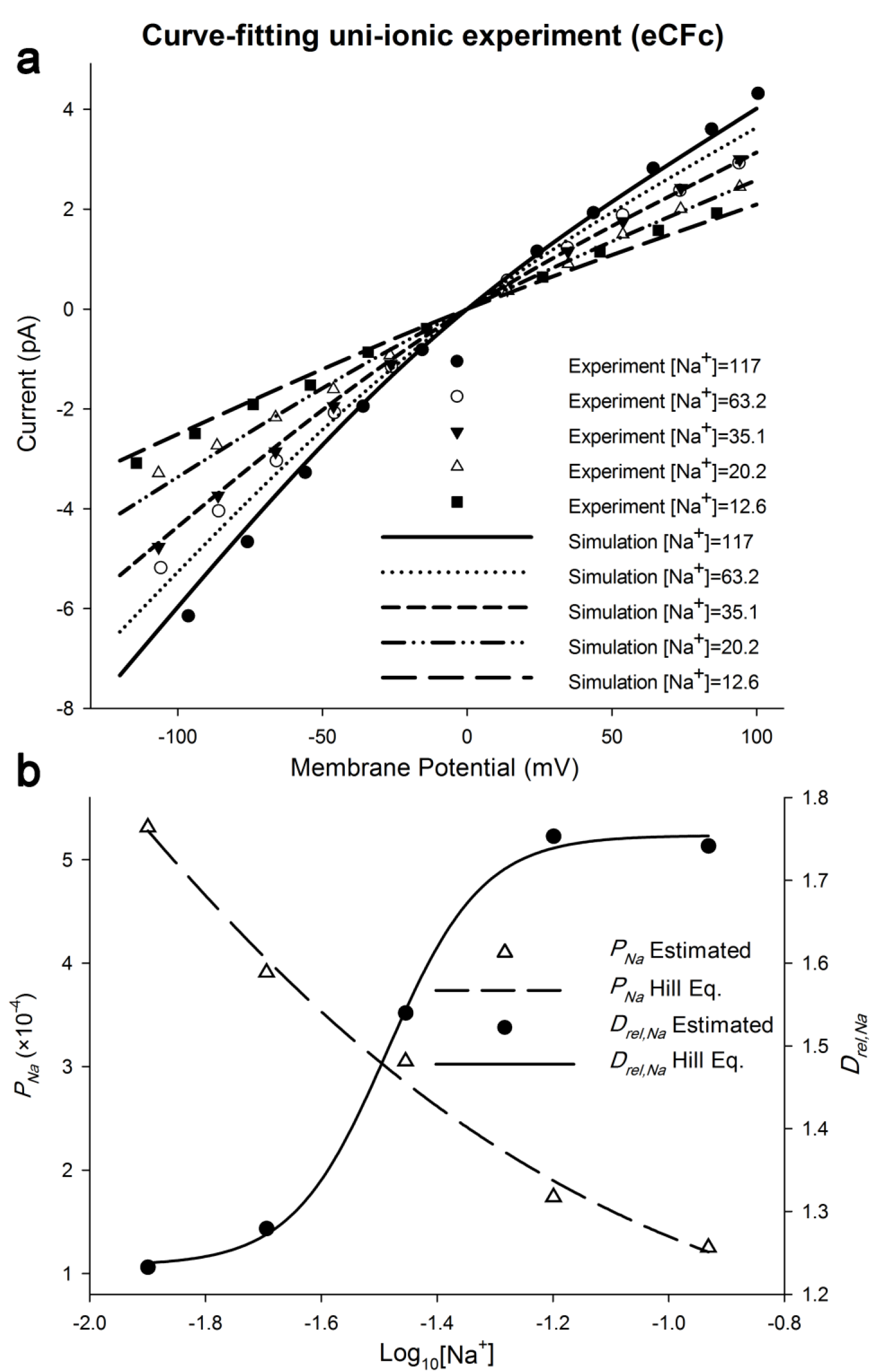
Hill equation models the concentration-dependence of eCFc equation parameters. The eCFc equation parameters are estimated by fitting it to the experimentally recorded single channel NMDAR current, in the uni-ionic experiment. NMDAR current is recorded at different holding potentials and symmetric sodium solution concentrations. (a) shows the curve-fitting accuracy. Na^+^ activities are reported in mM/liter. Note that the inward current compared to the outward current is about two times larger in higher concentrations. (b) shows the estimated parameters and their estimates by Hill equation.

In our previous study, we have introduced a fast Hodgkin and Huxley type model of NMDAR voltage-dependent gating (Moradi et al. 2012). In this study, we introduce an electrodiffusion model of NMDAR permeability that is also able to unify the permeability models of Ohmic and non-Ohmic channels. Starting from the Nernst-Planck electrodiffusion equation, we have reached to two extended forms of the constant field equation that are more precisely applicable to the biological channels. Our aim is to show that our constant field equations in conjunction with Hill equation are able to simulate the ion permeation of the NMDAR channel at different ionic concentrations and membrane potentials.

## 2. Material and Methods

### 2.1 The experimental data

We used the experimental data of Iino et al.(Iino et al. 1997) in which the recombinant ζ1 ε2 NMDARs have been used to obtain the single channel *I-V* curves of these receptors in the symmetric solutions of Na^+^ (the uni-ionic experiment), and in the asymmetric solutions of Na^+^ and Ca^2+^ in which Ca^2+^ was present in the extracellular solution only (the bi-ionic experiment).

### 2.2 The analyses

All the modeling and the optimization parts of this study were done in the Python programming language using OpenOpt, SciPy and NumPy packages. We used the least-square method to calculate the errors of our model with respect to the experimental data. The “particle swarm” algorithm was used to find model’s parameters by minimizing the error (Vaz and Vicente 2007).

### 2.3 Calculations

#### 2.3.1 The derivation of “general Constant Field current” equation

The oCFc equation was first introduced to describe the ion permeation in the bio-membranes (Hille 2001). In the “constant field theory”, it is assumed that a small and rate limiting fraction of ions in the solution are able to enter the channel’s mouth. To determine this fraction’s concentration (*c*s) on each side of the channel, the activity of the permeant ion *S* on that side of the channel ([*S*]) is multiplied by the “partition coefficient” (*β*) of that side. The ionic activity, which is a temperature, pressure, and concentration-dependent parameter, shows the “effective ionic concentration” in the solutions. This parameter is not only a function of its own concentration but also of the other ionic species.

Here, we introduce a more generalized version of the oCFc equation that is more compatible with ionic channels. The ionic movements inside the channel are under the influence of the electric field and the concentration gradient across the channel. If we consider the channel’s extension in the membrane along the X axis, where *x* represents the location (please see the Fig. 5), it is possible to model this condition with a modified version of Nernst-Planck electrodiffusion equation, in which the diffusion coefficient is a function of location within the channel:

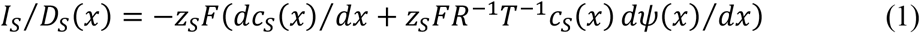

where, *I*_S_ is the amount of current density vector in direction *x* (A/cm^2^). *ψ*(*x*) is the local electrical potential inside the channel (in Volts). *c*_s_(*x*) is the local ionic concentration inside the channel (in M/cm^3^). *D_s_*(*x*) is the diffusion coefficient of ion S within the channel (in cm^2^/second) and *z*_S_ is its valance. The *F, R* and *T* are the Faraday’s constant, the gas constant and absolute temperature, respectively. The Nernst-Planck equation is a “non-exact differential equation” that can be solved by applying an “integrating factor”. The appropriate integrating factor *f*(*x*) is:

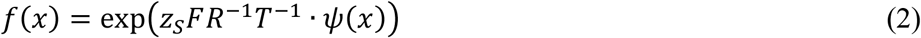

Multiplying the *f*(*x*) by the both sides of the Eq. 1, we would have:

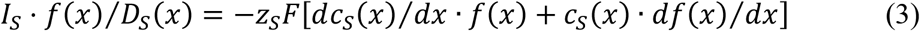

Integrating both sides, and simplifying the left side of the equation based on the product rule, our equation will be:

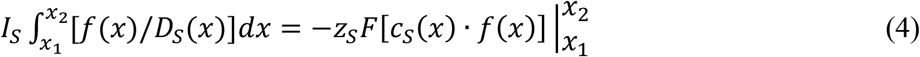

It is commonly accepted that the biological membranes behave like a capacitor. The electric filed between the two plates of a capacitor, *E*, is a constant. Consequently, the local electric potential changes linearly inside the membrane (*ψ*(*x*)=*E.x*). According to a recent molecular dynamic simulation study, it is a proper approximation to assume that the *ψ*(*x*), also changes linearly inside the transmembrane proteins (Gumbart et al. 2012). Membrane potential, *V*, is the potential difference between the intracellular and the extracellular parts of the membrane, where the potential in the extracellular part of the cell conventionally assumed to be zero. Therefore, if we assume that the electric potential inside the cell is zero (as is in the original solution of oCFc equation), so that the electric potential is –*V* in the extracellular fluid (Fig. 1), then *ψ*(*x*)=-*V.x/l*, where *l* is the channel’s length and *x* is changing from 0 (the inner mouth) to *l* (the outer mouth). If we substitute this Eq. to the Eq. 2 it will be:

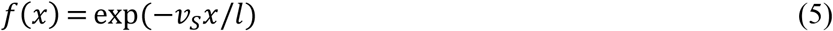

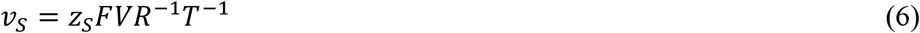

where, *v_S_* is a dimensionless parameter that summarizes all the constants in Eq. 1. Since we know the value of *c_s_* on each side of the channel, integrating Eq. 4 along the channel, and substituting Eq. 5 in it, we would have:

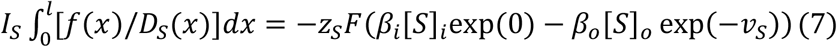

where, *β_i_* and *β_o_* are the partition coefficients of the inner and the outer parts of the channel, and [*S*]_*i*_ and [*S*]_*o*_ (in M/cm^3^) are the intracellular and the extracellular activity of the ion S in the solution, respectively.

To reach our “general Constant Field” current (gCFc) equation, we solve the left side of Eq. 7 assuming that a *δ* factor (0 < *δ* < 1) is dividing the channel into inner and outer parts, so that the diffusion coefficients, *D_s_*(*x*), of the inner and the outer parts have constant values *D_i,s_* and *D_o,s_* (see Fig. 1), respectively. Therefore, the integral in Eq. 7 would be the sum of two simpler integrals that we show it in the denominator of Eq. 8:

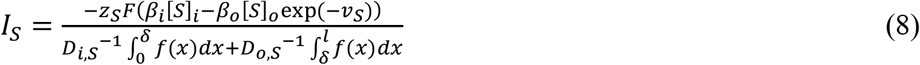

Substituting *f*(*x*) from Eq. 5 in this equation, calculating the integrals, and moving the negative sign from the nominator to the denominator, we would have:

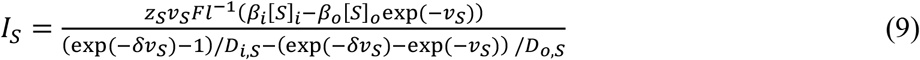

#### 2.3.2 The “extended Constant Field current” equation

In order to reach a simpler equation, the channel’s permeability (*P_s_*) should be defined. In the constant field theory, *P_s_* is an experimentally defined parameter that is introduced to calculate the flux of a permeant nonionic substance based on its activity difference in the intracellular and the extracellular solutions(Hille 2001). If we consider that the partition coefficients are equal (*β_i,s_* = *β_o,s_* = *β_s_*), then we would have:

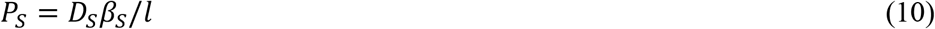

where, *D_s_* is the average diffusion coefficient within the channel. If we consider that *δ* = 0.5, then *D_s_* = (*D_i,s_* + *D_o,s_*)/2. In this particular case, it is possible to calculate *l* based on Eq. (10):

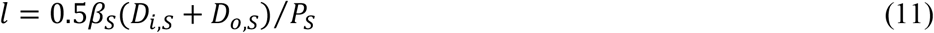

Solving Eq. 9 for *δ*=0.5, and substituting *l* from Eq. 11, we reach our “extended Constant Field” current (eCFc) equation that has fewer parameters than those of the gCFc equation:

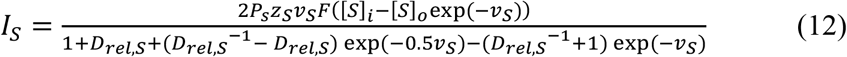

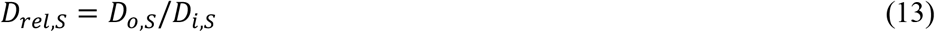

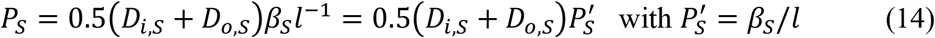

This equation takes into account the relative diffusion coefficient, *D_rei,S_*, of the inner and the outer parts of the channel. As a proof of the accuracy of our calculations, it is possible to show that this equation simplifies to the respected oCFc equation, when *D_o,s_* is equal to *D_i,s_*.

## 3. Results

To show our current equations’ ability to solve a real world problem, we fit them to the experimentally recorded NMDAR currents. To this end, we have used the experimental data of Iino et al., because in that study the *I-V* curve is obtained from the main open-state of the NMDARs in the single channel mode. Their approach simplifies the problem by bypassing the gating-related complexities. The experimental data of Iino et al. consist of two parts: 1) the uni-ionic experiment in which the *I-V* curves of NMDARs are obtained in the symmetric concentrations of sodium ions at different concentrations, and 2) the bi-ionic experiment in which the *I-V* curve is obtained in the asymmetric concentrations of sodium and calcium ions in which calcium ions were present in the extracellular solution only.

Our model is based on our general and extended constant field current equations, (see the calculations section). We derived the gCFc equation, Eq. 9, with the idea in mind that the selectivity filter of a channel, which is located in the relative position of *δ*, divides it into intracellular and extracellular parts. Each side of the channel has its own partition (*β_i,s_* and *β_o,s_*) and diffusion coefficients (*D_i,s_* and *D_o,s_*), so that the ion binding-sites, by changing the diffusion coefficients inside the channel, are able to modify the diffusion potential as well as the electric potential. The diffusion coefficient shows the ease with which ions can pass through the channel, while the partition coefficient determines the portion of ions in the solution that reach the channel mouth.

The eCFc equation, Eq. 12, results from the gCFc equation when we assume that the selectivity filter is located in the middle of the channel (*δ*=0.5), and the *β_i,S_* is equal to *β_o,S_*. The gCFc equation, directly calculates the ionic flux based on diffusion coefficients and partition coefficients but the eCFc equation, like the oCFc equation, calculates the ionic flux based on the ionic permeability (*P_S_*). Compared to oCFc equation, the eCFc equation has an additional parameter, named *D_rel,S_*, which shows the asymmetry of diffusion coefficient. The eCFc equation simplifies to the oCFc equation whenever there is no asymmetry in the ionic diffusion coefficients.

Our modeling study consists of two sections. In the first section, we analyze the current in the uni-ionic experiment. We model the sodium current (*I_Na_*) with eCFc equation. The appropriate values of *P_Na_* and *D_rel,Na_* at each concentration of ions are estimated by fitting the model to the experimental data. Then, the best function that is able to describe the concentration dependence of these parameters is found. In the second section, we make a pool of all the experimental data consisting of uni-ionic and bi-ionic experiments and analyze it as a whole. Our model consists of a system of equations including a pair of gCFc equations (one for *I_Na_* and one for *I_Ca_*) and a set of empirical equations that estimate the parameters of these current equations at each concentration of ions.

### 3.1 The uni-ionic experiment

In contrast to Ohm’s law or the oCFc equation, the *I-V* curve of NMDAR channel is not linear in the symmetrical solutions of sodium ions. The experimental findings show that the inward NMDAR current is more prominent than the outward, but *E*_rev_ is about 0 mV, as predicted by the Nernst equations (Iino et al. 1997). It seems, the *β_o,S_* = *β_i,S_* = *β_S_* according to the Nernst equation and the constant field assumptions (see the discussion section). If we also assumed that *δ* = 0.5, eCFc equation would be a valid model for NMDARs.

Giving a value greater than one for the *D_rel,Na_*, and an appropriate value for the permeability of sodium ions (*P_Na_*), our eCFc equation is able to simulate this phenomenon at different concentrations of sodium ions (Fig. 2a). Our simulations show that by increasing the Na^+^ concentration in the solution, *P_Na_* is reduced and *D_rel,Na_* is increased (Fig. 2b). Therefore, it seems that the sodium ions have a blocking effect on the permeability, which seems to be more powerful in the inner part of the channel. We realized that the best equation that is able to describe the concentration dependence of these parameters is the general form of the Hill equation, which is a sigmoid shape function:

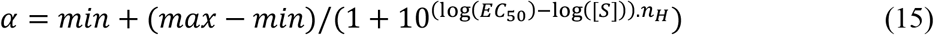

where, *α* is *P_Na_* (or *D_rel,Na_*), the *min* and *max* determine the minimum and the maximum of the function, respectively; *EC50* is the “half-maximal effective activity”, [*S*] is the activity of ion (here the sodium ions) and *n_H_* is the “Hill coefficient” that determines the slope of the function. In previous studies, this equation has been used to model a substance binding to its binding site; for a review see (Goutelle et al. 2008).

### 3.2 The pooled experimental data

To make a model that is able to predict the current at different ionic concentrations, we made a system of equations that estimates the eCFc parameters at different ionic concentrations. Parameters of this model were found by fitting it to a pool of all the experimental data. The *P′_S_*, *D_o,S_* and *D_i,S_* were estimated directly with a bound constrained optimization method. Eqs. 13 and 14 used to calculate eCFc parameters. Our model is based on some logical assumptions that simplify the problem:

1. The net current results from the sum of *I_Na_* and *I_Ca_*. These currents are modeled with separate eCFc equations. Since the increasing concentrations of calcium ions decrease the inward current in the Iino et al. experiment, we assume that the external calcium is able to reduce the *D_o,Na_* so that the net inward current becomes smaller than that of the uni-ionic experiment:

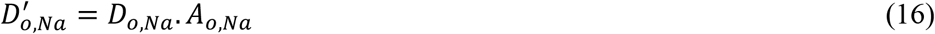

where, 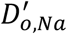 is the diffusion coefficient of sodium ions in the outer part of the channel, and in the presence of calcium ions, *D_o,Na_* is the sodium diffusion coefficient when there is no calcium in the solution, and *A_o,Na_* is the calcium adjustment ratio of *D_o,Na_*, which is a function of external calcium concentration and was limited to a value between zero and one. When there is no calcium, there should be no calcium block and therefore the maximum of the calcium adjustment ratio was fixed to be one. The minimum adjustment ratio, however, was estimated by curve-fitting.
2. Based on our simulations with the eCFc equation in the previous section, it seems that the Hill equation can explain the concentration dependence of parameters. Therefore, *D_i,Na_, D_o,Na_, D_o,Ca_* and *A_o,Na_* were modeled with the Hill equation. The *D_i,Na_* and *D_o,Na_* were a function of sodium activity in the internal and the external solutions, respectively. The *_Do,Ca_* and *A_o,Na_* were dependent on calcium activity in the external solutions.
3. The 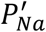 and 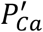 were constrained to be between zero and 10^-3^, in order to find comparable results with that of uni-ionic experiment. The optimization results of this model are summarized in table 1 and Fig. 3. Note that all the diffusion coefficients are decreasing with increasing ionic concentrations.

**Fig. 3.**
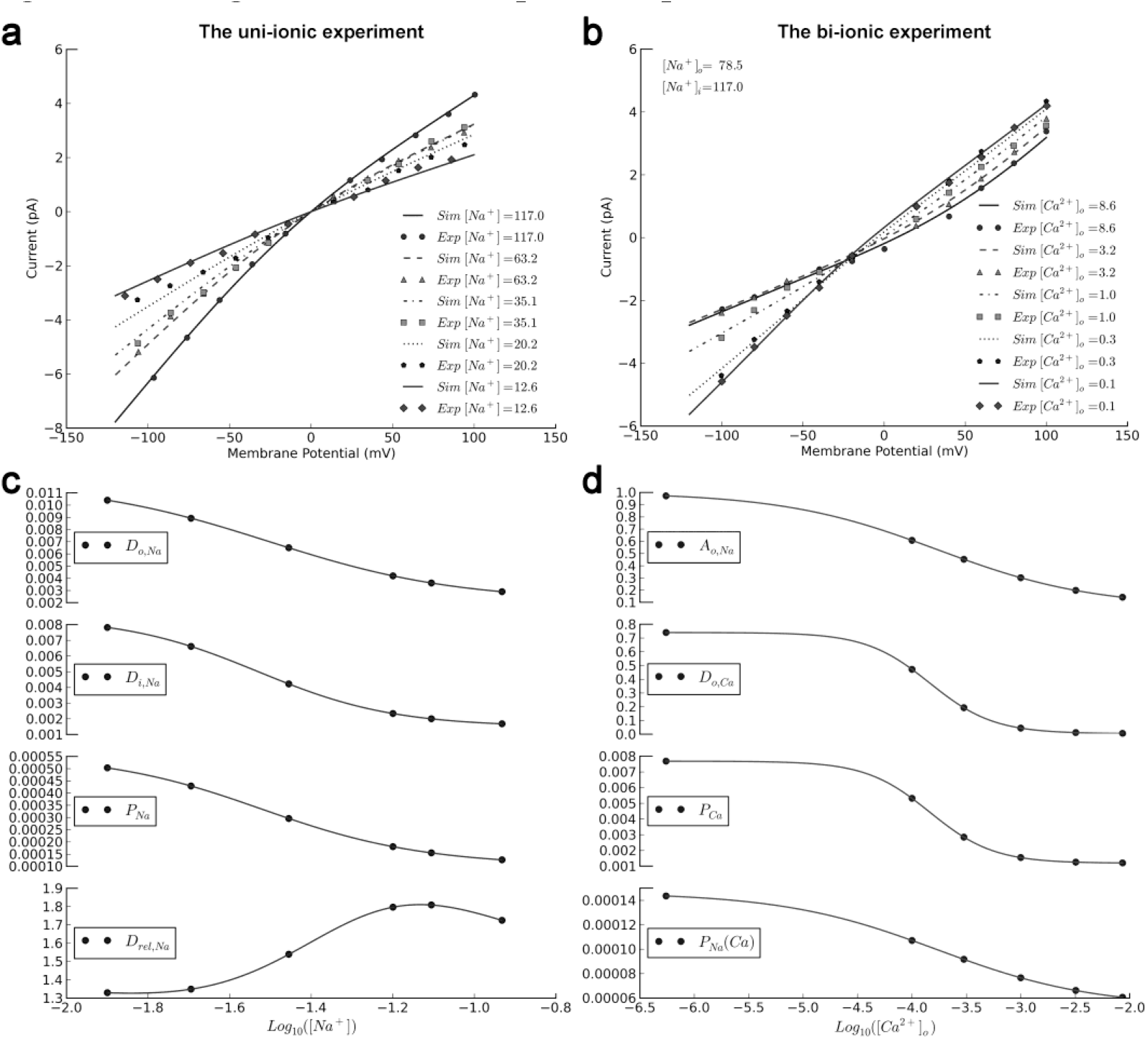
The fitting our model to the pooled experimental data. The experimental data consists of the NMDAR *I-V* curve in the symmetric uni-ionic Na+ solutions (the uni-ionic experiment), and the asymmetric Na^+^ and Ca^2^+ solutions (the biionic experiment). In the bi-ionic experiment, the [Na^+^] has a fixed value, while the [Ca^2+^]_o_ increases. *I_Na_* and *I_Ca_* modeled with two gCFc equations, and their parameters are calculated with Hill equation. (a, b) show the *I-V* curve in different concentrations, and the curve-fitting accuracy. Ionic activities are in mM/liter. (c) shows the concentration-dependence of *D_i,Na_* and *D_o,Na_* in the uni-ionic experiment. The eCFc parameters are calculated based on the estimated values of *D_i,Na_, D_o,Na_* and *P′_Na_* (d) shows the concentration-dependence of *D_o,Ca_* and *A_o,Na_*, in the bi-ionic experiments. *P_Na_* after Ca^2+^-block (*P_Na_*(*Ca*)) and the other eCFc parameters in the bi-ionic experiment are also calculated based on the estimated values of *D_o,Ca_, D_i,Ca_, A_o,Na_, P′_Na_*, and *P′_Ca_*.

**Table 1.** The parameters of the system of equations that models NMDAR.

| **Parameters** | ***min*** | ***max*** | ***EC*<sub>50</sub>** | ***n<sub>H</sub>*** | ***Variable (mM/liter)*** |
| --- | --- | --- | --- | --- | --- |
| ***A*<sub>o,Na</sub>** | 6.165e-02 | 1 | 1.721e-04 | -6.074e-01 | [Ca<sup>2+</sup>]<sub>o</sub> |
| ***D*<sub>o,Na</sub>** | 2.155e-03 | 1.187e-02 | 3.134e-02 | -1.886e+00 | [Na<sup>+</sup>]<sub>o</sub> |
| ***D*<sub>i,Na</sub>** | 1.499e-03 | 8.501e-03 | 2.955e-02 | -2.623e+00 | [Na<sup>+</sup>]<sub>i</sub> |
| ***D*<sub>o,Ca</sub>** | 4.500e-03 | 7.404e-01 | 1.456e-04 | -1.478e+00 | [Ca<sup>2+</sup>]<sub>o</sub> |
| ***P*'<sub>Na</sub>** | 5.521e-02 | - | - | - | - |
| ***P*'<sub>Ca</sub>** | 1.765e-02 | - | - | - | - |
| ***D*<sub>i,Ca</sub>** | 1.301e-01 | - | - | - | - |

### 3.3 The NMDAR current analysis

Building a model that in mixed ionic conditions is able to dissociate the NMDAR current to its main ionic components was one of our main incentives in this study. For the bi-ionic experiment, the dissociated *I_Na_* and *I_Ca_*, the relative calcium current (*I_Ca_/I_Na_*), the fractional calcium current (*P_f_* = *I_Ca_*/(*I_Na_*+*I_Ca_*)×100), Erev and *P_Ca_/P_Na_* are plotted in Fig. 4. Note that at membrane potentials of about −25 mV, i.e. the region in which the degree of magnesium block is at its minimum, the calcium flux does not change significantly in the [Ca^2+^]_o_ range of 0.3 to 3.2. It is also worth noting that *I_Ca_*/*I_Na_* depends on the Erev of *I_Na_*, while the *P_f_* depends on the Erev of the total NMDAR current. Also, note that the *P_f_* and *I_Ca_/I_Na_* are voltage independent when the difference between the value of membrane potential and the Erev is large enough.

**Fig. 4.**
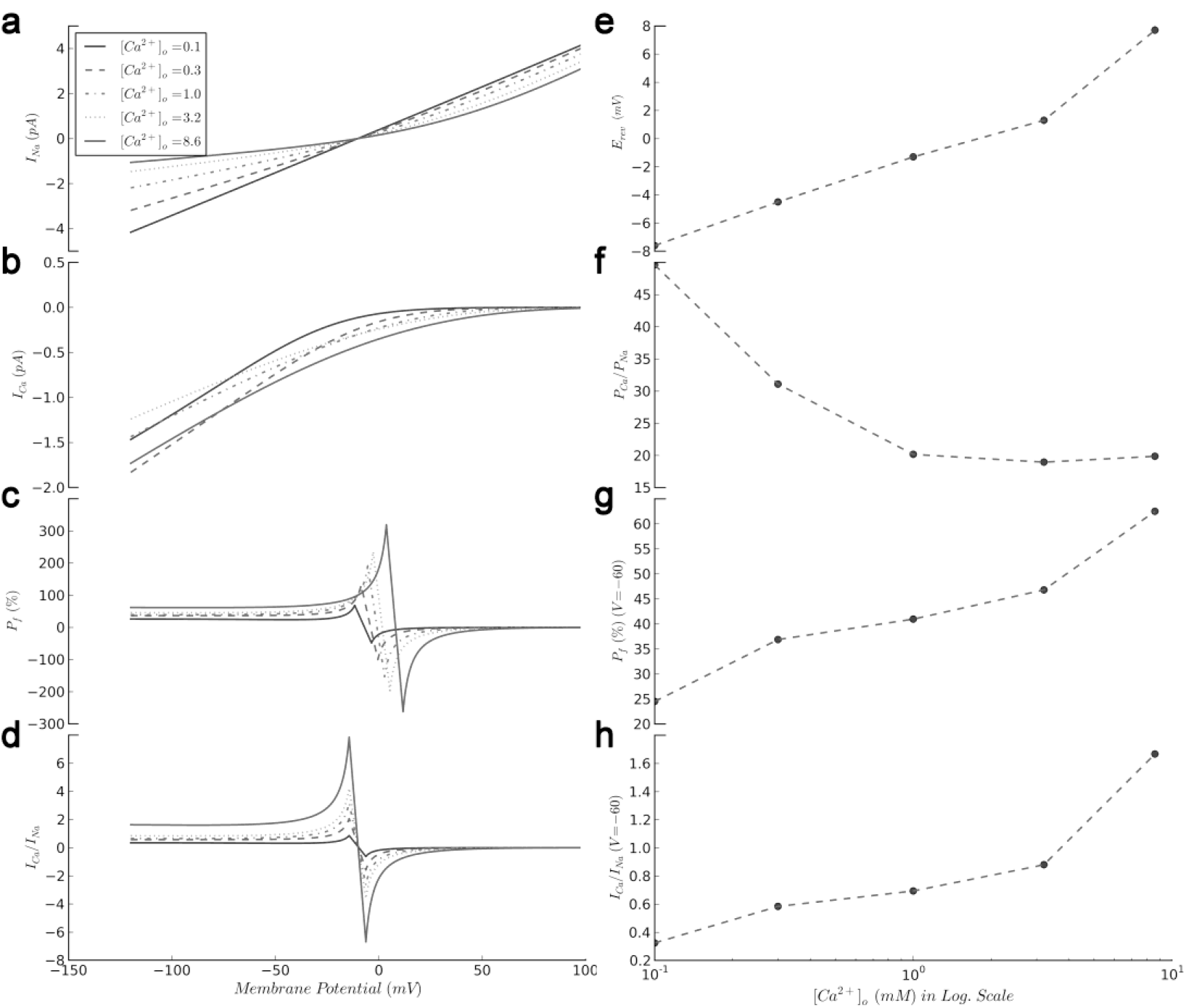
The current analysis in the bi-ionic experiment. Different ionic permeability indices are calculated based on our fitted model of bi-ionic experiment. (a) shows the sodium current, (b) shows the calcium current, (c) is the fractional calcium current, and (d) is the relative calcium to sodium current, all against the membrane potential. (e) is the reversal potential, (f) is the relative permeability of calcium to sodium, (g) is the fractional calcium current, and (h) is the relative calcium to sodium current, all against the calcium concentration.

## 4. Discussion

The oCFc equation’s validity is widely accepted among the physiologists, but it fails to model the non-Ohmic NMDAR current. In the Jahr and Stevens Ohmic model, this current is simulated by considering different conductances for the inward and the outward currents(Jahr and Stevens 1993). The oCFc equation is also able to separate the inward and the outward permeabilities:

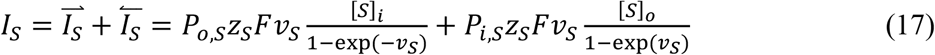

Although theoretically incorrect (please see the introduction section), in a computational model, we gave different values to *P_i,S_* and *P_o,S_* in order to build a model that is theoretically close to their model. Unlike their model, however, this equation does not need a transition function. Our simulations showed that neither this equation nor Jahr and Stevens’ model fits to the pooled experimental data of Iino et al. (results are not reported). For instance, such an oCFc equation is unable to predict the Erev at the uni-ionic experiment (≈ 0 mV).

The “macroscopic electrodiffusion rules” application to study “microscopic ion channels” has been a controversial topic. Since the oCFc equation cannot simulate the NMDAR current even with the above-mentioned modifications, we conclude that some of the assumptions underlying the oCFc equation should not be valid, at least in the NMDARs case. The oCFc equation is derived from the Nernst-Plank electrodiffusion equation, if certain conditions are assumed: 1) the membrane potential should change linearly inside the channel, 2) the inner and the outer partition coefficient should be equal, and 3) the diffusion coefficient should be constant within the channel.

A recent molecular dynamics simulation study of the membrane proteins shows that the linear function can be a reasonably acceptable model of the electric field inside a channel(Gumbart et al. 2012). In addition, all the significance of the constant field theory that it can predict the Nernst equation and the Ohm’s law, comes from the hypothesis that the electric potential changes linearly inside a membrane. To explain this linearity, maybe it is possible to assume that a binding site that can attract a positively charged ion should have negative charge that is able to attract positive charges until it is completely neutralized. Even if the binding-site charge were not neutralized completely, at the worst-case scenario, we would have an electric dipole of two ions located in the Y-Z plane (see Fig. 5) that is going to affect the whole membrane’s electric field, which includes a vast amount of ions on membrane. Because the channel diameter is smaller than its length, the impact of this dipole on the Y-Z plane (which affects diffusion coefficient via the frictional forces) would be larger than the X axis (which is the direction of the effective electric field). In addition, when the passing charge is getting close to the dipole, the potential energy of the ion would be stored but when the ion goes away, this potential energy would be released. Therefore, the net potential change due to the binding site (dipole) would be zero. Because of the dipole’s small atomic size compared with the channel’s large macromolecular-sized length, effect of this change in the electric potential can be negligible. Therefore, in our equations, it is reasonable to assume the membrane potential (but not the diffusion coefficient) a linear function.

**Fig. 5.**
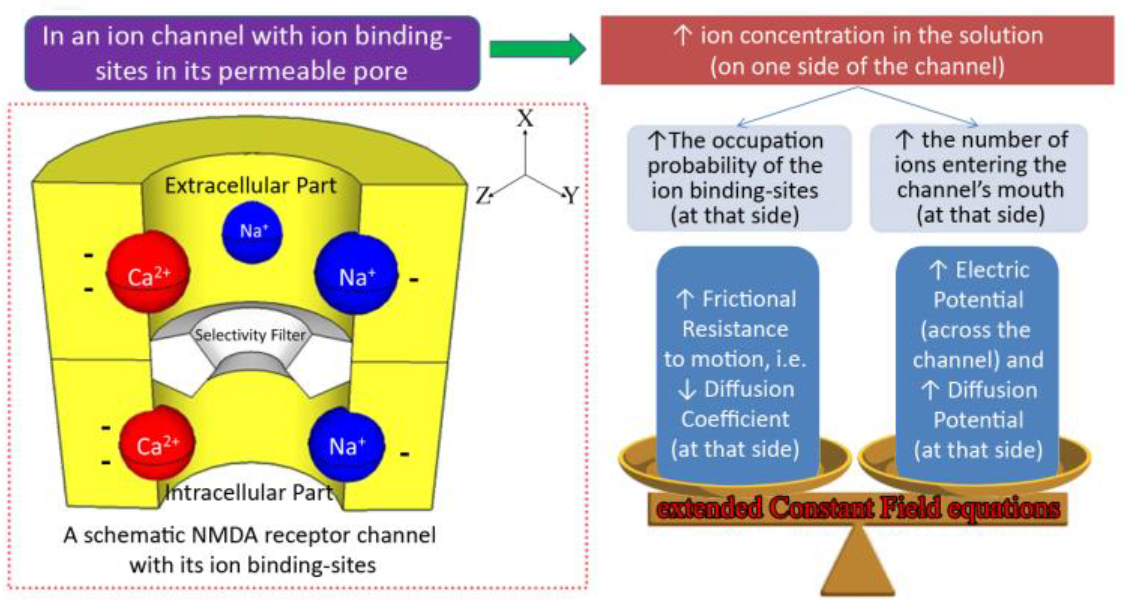
the NMDAR schematic model.

Historically speaking, the partition coefficient was introduced when researchers were trying to study the permeability of non-ionic substances through the biomembranes. The idea was that a substance’s permeability is determined by its solubility in membrane, and the partition coefficient represents this solvation degree. Lipid-bilayer can be a symmetric structure and therefore in the oCFc equation, the inner and the outer partition coefficients are considered equal. In our constant field channel model (gCFc), we did not find any reason to consider the inner and the outer partition coefficients equal.

Although in theory, membrane behaves like a capacitor and therefore there is no electric field outside the space between the two plates of this capacitor to affect the partition coefficients, it is possible that the asymmetric physical dimensions and/or the channel’s local surface electric charges affect these parameters. Therefore, we used different parameters for the inner and the outer partition coefficients in our gCFc equation. In the NMDAR’s case, however, *β_i,s_* should be equal to *β_o,s_* since the *E*_rev_, in agreement with the Nernst equation, was equal to zero in the symmetric uni-ionic solution. The Nernst equation predicts zero *E*_rev_ in the absence of electric and diffusion potentials. Since the Nernst equation can be derived from the oCFc equation (Hille 2001), the underlying assumptions of this equation (about the partition and diffusion coefficients) also apply to the Nernst equation. Considering the fact that the frictional forces are absent in the absence of driving forces, the partition coefficients are the sole determinants of Erev in this condition. Therefore, we kept *β_i,s_* =*β_o,s_* in our eCFc equation, in agreement with the previous studies (Sharma and Stevens 1996; Zarei and Dani 1994; Iino et al. 1997), and focused on the diffusion coefficient non-uniformity inside the channel. Diffusion coefficient is the inverse of frictional resistance to motion(Hille 2001). Thus, it should be a function of the channel’s dimensions, and the electrical forces between the permeating ions and the channel wall. Our simulations with the gCFc equation showed that the concentration-dependent change in the diffusion coefficient on the channel’s either side has a significant role in shaping the NMDARs’ *I-V* curve. We also showed that such changes obey the Hill equation, which has been used in the previous studies to model the binding of a molecule to its binding site. Consequently, the ion and binding sites interaction may reduce the ions permeation. The ion-binding-site existence inside the NMDARs is in agreement with the previous studies. Two binding-sites have been identified for monovalent cations in the channel’s extracellular part, and one binding site in its intracellular part, which are able to modify the channel’s permeability (Antonov et al. 1998; Zhu and Auerbach 2001b, 2001a). Similarly, in our model, both *D_o,Na_* and *D_i,Na_* decrease with increasing Na^+^ concentrations in the external and internal solution. When the Na+ concentration is low, the Na^+^ binding-probability is low on the channel’s both sides, and therefore, the *D_rel,Na_* gets close to 1. However, increasing the concentration, the *D_rel,Na_* gets close to 2, since these binding sites have different binding-probabilities.

In addition, two binding sites are identified for the divalent cations. One is located in the deeper parts of the channel, and the other one is a more external binding site that is called DRPEER(Sharma and Stevens 1996; Premkumar and Auerbach 1996; Watanabe et al. 2002). The concentration-dependent decrease in the *D_o,Ca_* and *A_o,Na_* shows that not only the Ca^2+^ but also the Na^+^ permeation is reduced by the Ca^2+^-bound binding sites. We have to use separate functions for the *D_o,Ca_* and *A_o,Na_*, probably because these ions have different diameters and electrical charges. Therefore, the Ca^2+^-bound binding sites reduce the calcium *D_o,Ca_* and *D_o,Na_*, differently. For a schematic of our model see Fig. 5.

### 4.1 The ionic permeability and the diffusion coefficient asymmetry

The *P_S_* is the common parameter of both eCFc and oCFc equations, which is introduced in previous studies to simplify the experimental study of substance permeation through biomembranes. In both of these equations, this parameter is determined by partition coefficient and average diffusion coefficient. In experimental studies, an Ohmic channel’s permeability is considered a constant that is concentration-independent. Simply, knowing the relative permeability to different ions, it is possible to model these channels with oCFc equation. The oCFc equation, however, neither models the nonlinear *I-V* curve of non-Ohmic channels, nor their concentration-dependent change in the permeability (D. Chen et al. 1997). Our model shows that the concentration-dependent change in the ionic diffusion coefficient inside the NMDAR channels not only explains the change in their permeability, but also their non-linear *I-V* curve (see Eqs. 13, 14 and Fig. 3c). We also show that not only Ca^2+^ (Iino et al. 1997; Jahr and Stevens 1993) but also Na^+^ reduces its own permeability in NMDARs. This finding is in agreement with a previous study where it is shown that Na^+^ blocks some organic ion’s permeation (Antonov et al. 1998).

In our model with gCFc equation, the ionic flux depends on three opposing factors: Two of them are the voltage and concentration-dependent mechanisms that increase the diffusional forces, and the third is the concentration-dependent reduction of the diffusion coefficient that reduces both of these forces. Ionic concentration is a powerful factor. For instance, when we used the *P_Na_* that was estimated by fitting eCFc (or oCFc) equation to the NMDAR’s current in [Na^+^]=12 mM symmetric solution in order to simulate the current in [Na^+^]=117 mM, the simulated current overshot the experimentally recorded values. Consequently, the *P_Na_* should be reduced by means of diffusion coefficient reduction to correct model estimations. Likewise, the *P_Ca_* is reduced by increasing [Ca^2+^]_o_, so that the Ca^2+^ flux remains an invariant value in a wide range of [Ca^2+^]_o_ (see *I_Ca_* at −-25 mV in Fig. 4b). The calcium-dependent reduction in *P_Na_* was in amounts that *P_f_* increased by increasing [Ca^2+^]_o_.

### 4.2 The permeability ratios

In the current literature, the *P_Ca_/P_Na_* indicates the relative amount of Ca^2+^ to Na^+^ flux. In some of the previous studies, the *P_Ca_/P_Na_* is calculated by oCFc equation (Jatzke et al. 2002; Lewis 1979). Our study, however, shows that these calculations are not valid at least for NMDARs, because the diffusion coefficient asymmetry is not considered in those calculations. According to our model, while the *P_Ca_/P_Na_* is decreasing with increasing [Ca^2+^]_o_, both *P_f_* and *I_Ca_/I_Na_* are increasing (see Fig. 4f-h). Therefore, *P_Ca_/P_Na_* is incompatible with the other indices.

Owing to its theory-independent calculation method, the *P_f_* value seems to be a better indicator of the Ca^2+^ flux amount. In the previous studies, measuring the charge that is carried by calcium ions then dividing it by the total charge that is carried by all ions, the *Pf* has been calculated experimentally. The total charge is estimated by integrating the current that passes through the current-clamp probe in Δ*t* milliseconds; meanwhile the calcium charge is estimated based on the change in [Ca^2+^]_i_ that is measured by fura-2 fluorescent dye. Indeed, the fura-2 dye, like most of the other measurement devices, is able to measure the free calcium activity. To convert the change in the calcium activity to the calcium charge that is carried through the channel, its value should be corrected for the calcium concentration by dividing it by the “calcium activity coefficient” in that particular experiment, also multiplied by the Ca^2+^ valence. However, without the abovementioned corrections and in the external calcium concentration of about 1.8 mM, Jatzke et al. have calculated *P_f_* and have reached a value of about 13.5% at −60 mV. Correcting it for the reported activity coefficient (=0.57) and for the ionic valence, we have reached a *P_f_* value of ≈47.4%. Multiplying the external calcium concentration (1.8 mM) by the activity coefficient (=0.57), the calcium activity has been around 1 mM in Jatzke et al. study. In our study and in the 1 mM calcium activity, we have reached a *P_f_* value of ≈41% at −60 mV (see Fig. 4g), which is comparable to the Jatzke et al. results. Consequently, for the first time in our study, we have built an electrodiffusion NMDAR model that reproduces theory independent experimental results.

In summary, in this study we have extended oCFc equation and proposed a new mechanism by which channels’ ion binding-sites can regulate the ionic flux. Our equations help researchers to analyze the mixed-ionic NMDAR current and possibly other non-Ohmic channels by formulating the relation between the current, voltage, ionic concentration and ionic binding-sites. Consequently, our equations may help computational modelers in order to model the NMDAR-related phenomena more precisely.

